# Multi-Criteria Evaluation of *Hibiscus mutabilis* Cultivars for Food Development Using Delphi and Analytic Hierarchy Process

**DOI:** 10.64898/2026.11.05.764492

**Authors:** Yaoyuwei Fan, Yuqing Ma, Xiaoqing Shi, Yongjie Feng, Yiqiong Liu, Hong Wang, Xi Wang

**Author notes:** Corresponding author: Xi Wang.

## Abstract

This study evaluated 11 *Hibiscus mutabilis* cultivars from Chengdu Botanical Garden to assess their quality for food use and identify promising cultivars. An evaluation framework was developed using the Delphi method to select indicators and the analytic hierarchy process (AHP) to determine their weights. Overall scores ranged from 2.5785 to 3.6043, and the cultivars were classified into three grades. Grade I included *Hibiscus mutabilis* ‘Bairi Huacai’, *Hibiscus mutabilis* ‘Chongbanbai’ and *Hibiscus mutabilis* ‘Jinqiusong’; Grade II included *Hibiscus mutabilis* ‘Jinqiuhong’, *Hibiscus mutabilis* ‘Zuifurong’, *Hibiscus mutabilis* ‘Danbanhong’, *Hibiscus mutabilis* ‘Jinxiuzi’, *Hibiscus mutabilis* ‘Jinbiyu’ and *Hibiscus mutabilis* ‘Caixia’; whereas Grade III included *Hibiscus mutabilis* ‘Mudanfen’ and *Hibiscus mutabilis* ‘Jinrui’. In summary, *Hibiscus mutabilis* ‘Bairi Huacai’, *Hibiscus mutabilis* ‘Chongbanbai’ and *Hibiscus mutabilis* ‘Jinqiusong’ are the preferred cultivars for food processing and development. The framework provides a basis for cultivar screening and further evaluation of *H. mutabilis* flowers for food use.

## Introduction

*Hibiscus mutabilis* L. is a deciduous shrub or small tree belonging to the genus *Hibiscus* in the family Malvaceae[1]. Widely cultivated as an ornamental plant in China, *H. mutabilis* is valued for its diverse flower colours and forms, long flowering period, and adaptability to different growing conditions[2–4]. Recent efforts in germplasm collection and cultivar development have expanded the resources available for the diversified use of *H. mutabilis*[5–7]. Cultivars differ in flower colour[8], morphology[9], and regional adaptability[10], providing a germplasm foundation for their diversified utilisation. In addition to ornamental uses, the flowers of *Hibiscus* syriacus and its closely related species, as distinctive plant resources, also have a history of traditional culinary use[11–13]. *Hibiscus* flowers are used in tea and various culinary preparations[14]. Growing interest in plant-based food ingredients has prompted research into the nutritional and bioactive components of *H. mutabilis* flowers, including amino acids, mineral elements, and flavonoids[15–17].

Currently, relevant research has primarily focused on the ornamental characteristics[17], germplasm resources[5], cultivation adaptability[18] and medicinal value of *Hibiscus*[16]. However, comparative assessments of cultivars for food use remain limited, and a comprehensive evaluation framework is still needed[19]. Therefore, this study utilised *Hibiscus* flowers from 11 cultivars as the test material, employing the Delphi method to screen and select evaluation indicators[20–22], and combined this with the Analytic Hierarchy Process (AHP)[23] to construct a comprehensive evaluation system for the edible quality of *H. mutabilis*. This system was used to assign comprehensive scores, rank and classify the different cultivars, with the aim of providing a reference for the development of superior *Hibiscus* cultivars and their subsequent utilisation in food processing.

## Materials and Methods

### Plant Materials

Eleven *H. mutabilis* L. cultivars were collected from the research nursery of Chengdu Botanical Garden: *Hibiscus mutabilis* ‘Jinxiuzi’, *Hibiscus mutabilis* ‘Jinqiusong’, *Hibiscus mutabilis* ‘Chongbanbai’, *Hibiscus mutabilis* ‘Caixia’, *Hibiscus mutabilis* ‘Jinqiuhong’, *Hibiscus mutabilis* ‘Zuifurong’, *Hibiscus mutabilis* ‘Jinbiyu’, *Hibiscus mutabilis* ‘Bairi Huacai’, *Hibiscus mutabilis* ‘Jinrui’, *Hibiscus mutabilis* ‘Danbanhong’ and *Hibiscus mutabilis* ‘Mudanfen’ (Fig 1)(cultivars are referred to by cultivar name only in subsequent sections). Among these, *Hibiscus mutabilis* were developed by Chengdu Botanical Garden through its own breeding program. Fully opened flowers, intact and free of mechanical damage or visible pests and diseases, were sampled in mid-October 2025. For each cultivar, three plants were sampled per cultivar. Thirteen flowers from each plant were pooled to form one biological replicate, yielding three biological replicates per cultivar.

**Fig 1.**
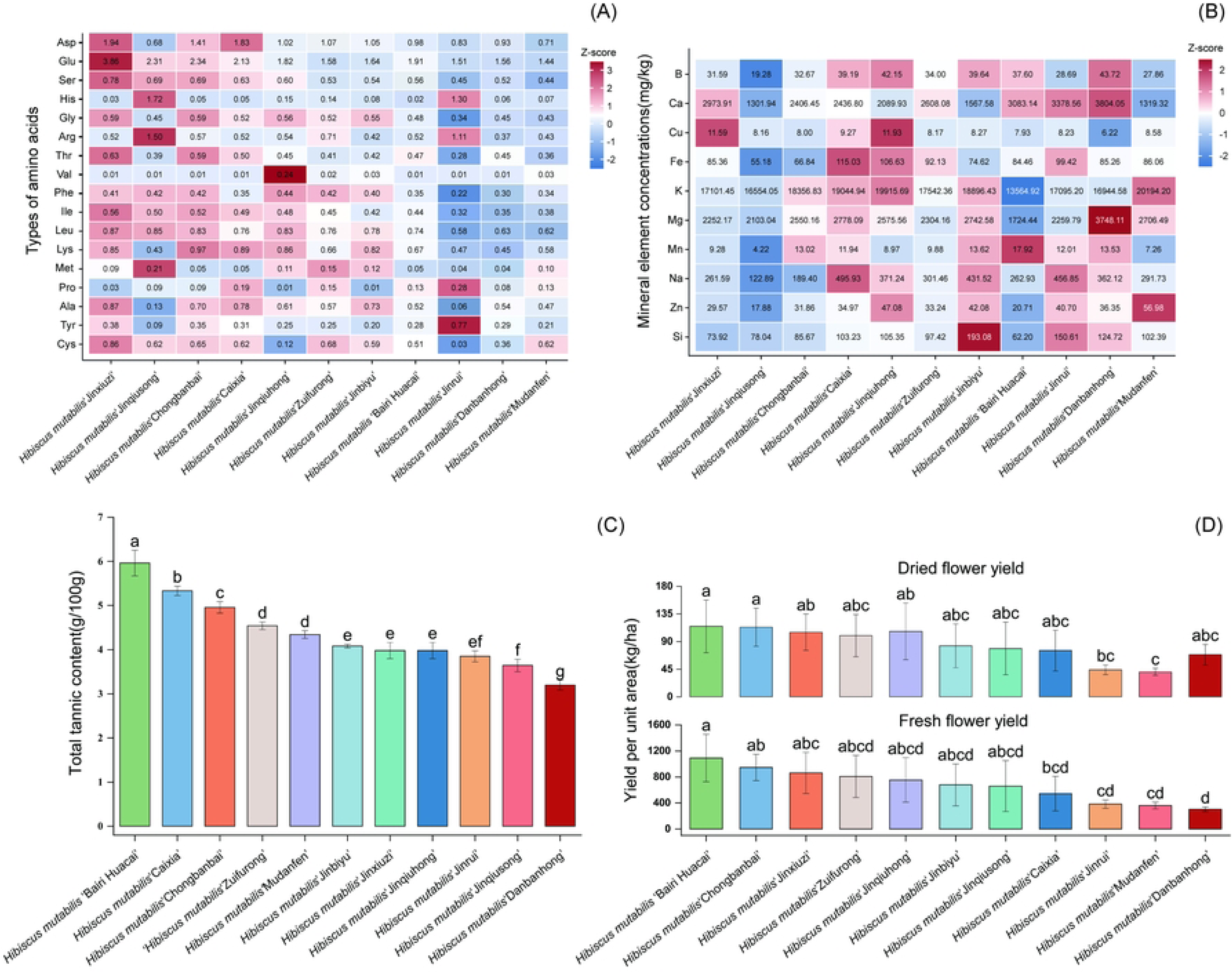
11 cultivars of *Hibiscus mutabilis*

### Screening of Evaluation Indicators

#### Construction of the initial evaluation index system

Based on the requirements for the edible quality of *H. mutabilis* flowers and relevant literature, and taking into account food safety, nutrition and active substances, sensory quality, and processing adaptability, an initial evaluation index system for the edible quality of *H. mutabilis* flowers was constructed. The initial indicators covered four aspects—safety, nutrition and active substances, sensory quality, and storage and adaptability—comprising 19 indicators in total (Table 1).

**Table 1.** Initial Evaluation Index System.

| Criterion layer | Specific indicator |
|---|---|
| Safety | 1. Heavy metal contamination risk |
|  | 2. Microbiological contamination risk |
|  | 3. Allergenicity |
|  | 4. gossypol-related toxicity |
|  | 5. Exogenous pollutant barrier capacity |
| Nutrition and active substances | 6. Total flavonoid content |
|  | 7. Soluble sugar content |
|  | 8. Total amino acid content |
|  | 9. Vitamin C content |
|  | 10. Total mineral element content |
| Sensory quality | 11. Fiber coarseness |
|  | 12. Lingering astringency |
|  | 13. Herbal aroma recognizability |
|  | 14. Mouthfeel smoothness |
|  | 15. Fading rate |
|  | 16. Appearance and color |
| Storage and adaptability | 17. Storability |
|  | 18. Gel-forming capacity |
|  | 19. Color stability |

#### Delphi expert consultation

This study employed the Delphi method to screen and optimise the initial evaluation indicators. An expert consultation questionnaire was designed based on the initially constructed 19 indicators, and experts in horticulture, plant resources, food safety, food processing, and *H. mutabilis*-related research were invited for evaluation. The questionnaire mainly covered three aspects: (i) A 10-point scale was used to assess the suitability of each indicator for evaluating the edible quality of *Hibiscus* flowers; (ii) Seek experts’ views on the addition, deletion, merging or amendment of indicators; (iii) Assess the relative importance of the four criteria levels: safety, nutritional and bioactive compounds, sensory quality, and shelf life and adaptability—using a five-level scale: unimportant, less important, moderately important, important, and very important.

In the first round, a total of 32 expert consultation questionnaires were distributed and 32 were returned, of which 28 were valid, representing a valid response rate of 87.5%. The participating experts were primarily from the fields of horticulture, botany, food safety, food processing and related research areas, providing good professional representation. Based on the experts’ scores, the mean, standard deviation and coefficient of variation for the importance scores of each indicator were calculated, and the indicators were adjusted in light of the experts’ written comments. The selection of indicators was primarily based on their importance, the degree of consensus among experts, their measurability and their practical application value. Indicators that were difficult to quantify, involved high testing costs or had a weak correlation with the evaluation of edible quality were either deleted or merged.

Based on the results of the first round of expert consultation, adjustments have been made to the initial evaluation indicator system. In the safety layer, ‘sensitisation’ and ‘ability to block exogenous contaminants’ were removed from the safety indicators; in the nutritional and active substance layer, ‘soluble sugar content’ and ‘vitamin C content’ were removed from the nutrition and active ingredients indicators; in the sensory quality layer, ‘herbal aroma recognisability’, ‘colour fading rate’ and ‘appearance and colour’ were removed from the sensory quality indicators; and in the shelf-life and adaptability layer, ‘gel-forming ability’ and ‘colour stability’ were removed from the shelf-life and adaptability indicators. To better assess the suitability of *H. mutabilis* flowers as raw materials for food development, yield per unit area and drying yield were incorporated into the revised evaluation framework. After the first round of revisions, the number of evaluation indicators was reduced from 19 to 12, and the revised indicator system was established for the second round of expert consultation.To enhance the alignment of the evaluation system with the objectives of food development and utilisation, the original criterion ‘storage stability and adaptability’ has been revised to ‘raw material supply characteristics’.To strengthen the link with food development and utilization, the original criterion of ‘storage and adaptability’ was adjusted ‘raw material supply characteristics’. After two rounds of expert consultation and revision, this study established an evaluation framework comprising four criteria—safety, nutritional and active substance, sensory quality, and raw material supply characteristics—and 12 evaluation indicators, as presented in Table 2.In particular, safety assesses the basic eligibility of *H. mutabilis* as food; nutritional and active substance establish its core nutritional and functional value; sensory quality determines consumer acceptability; and raw material supply characteristics evaluate the species’ capacity for a stable supply.

**Table 2.** Final Evaluation Indexes.

| **Criterion layer** | **Specific indicator** |
|---|---|
| Safety | 1. Heavy metal contamination risk |
|  | 2. Microbiological contamination risk |
|  | 3. Gossypol-related toxicity risk |
| Nutrition and active substances | 4. Total flavonoid content |
|  | 5. Total amino acid content |
|  | 6. Total mineral element content |
| Sensory quality | 7. Fiber coarseness |
|  | 8. Lingering astringency |
|  | 9. Mouthfeel smoothness |
| Raw material supply characteristics | 10. Yield per unit area |
|  | 11. Drying yield |
|  | 12. Storability |

### Construction of the AHP-Based Comprehensive Evaluation System and Weight Determination

The Analytic Hierarchy Process (AHP) was used to develop a comprehensive evaluation system for the edible quality of *H. mutabilis* flowers. The system comprises four hierarchical layers: the goal layer (A), the criterion layer (C), the indicator layer (P) and bottom layer (D).The goal layer provides a comprehensive evaluation of the edible quality of *H. mutabilis*; the criterion layer comprises safety (C1), nutritional and active substance (C2), sensory quality (C3), and raw material supply characteristics (C4); the indicator layer includes 12 specific evaluation indicators; and the bottom layer covers the 11 *H. mutabilis* cultivars under evaluation. The detailed indicator framework is presented in Table 3. Specifically, endogenous toxin risk is evaluated via free gossypol content, and Lingering astringency is evaluated via total tannin content.

**Table 3.** Evaluation model framework.

| Goal layer (A) | Criterion layer (C) | Indicator layer (P) | Bottom layer (D) |
|---|---|---|---|
| Comprehensive evaluation of the food development potential of *H. mutabilis* flowers | C1 Safety | P1 Heavy metal | D1–D11 *H. mutabilis* cultivars to be evaluated |
|  |  | P2 Microbiological |  |
|  |  | P3 Gossypol-related |  |
|  | C2 Nutritional and active substance | P4 Total flavonoid |  |
|  |  | P5 Total amino acid |  |
|  |  | P6 Total mineral |  |
|  | C3 Sensory quality | P7 Fiber coarseness |  |
|  |  | P8 Lingering |  |
|  |  | P9 Mouthfeel |  |
|  | C4 Raw material supply characteristics | P10 Yield per unit |  |
|  |  | P11 Drying yield |  |
|  |  | P12 Storability |  |

Based on expert consultation results, a 1–9 scale was used for pairwise comparisons of indicators at the same hierarchical level to construct the decision matrix. SPSSAU was employed to calculate the maximum eigenvalue, weight vector, and consistency index for each matrix. The consistency ratio (CR) was then computed to assess matrix consistency, as expressed in the following formula:

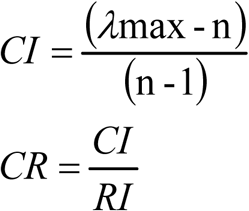

where λmax is the maximum eigenvalue of the judgment matrix, n is the matrix order, and RI is the average random consistency index. When CR < 0.10, this indicates that the matrix exhibits satisfactory consistency (Table 4).

**Table 4.** A-C Consistency Test of Judgment Matrix.

| **Layer** | **Matrix** | **Column criterion order** | **Row indicator** | **Judgment value** | **Weight W** | **λmax** | **CI** | **CR** | **Consistency** |
|---|---|---|---|---|---|---|---|---|---|
| Goal layer-Criterion layer | A-C | C1,C2,C3,C4 | C1 | 1,2,3,2 | 0.4168 | 4.071 | 0.024 | 0.027 | Pass |
| Goal layer-Criterion layer | A-C | C1,C2,C3,C4 | C2 | 1/2,1,2,2 | 0.2695 | 4.071 | 0.024 | 0.027 | Pass |
| Goal layer-Criterion layer | A-C | C1,C2,C3,C4 | C3 | 1/3,1/2,1,1/2 | 0.1209 | 4.071 | 0.024 | 0.027 | Pass |
| Goal layer-Criterion layer | A-C | C1,C2,C3,C4 | C4 | 1/2,1/2,2,1 | 0.1928 | 4.071 | 0.024 | 0.027 | Pass |
| C1 sub-criteria layer | C1-P | P1,P2,P3 | P1 | 1,5,3 | 0.6333 | 3.039 | 0.019 | 0.037 | Pass |
| C1 sub-criteria layer | C1-P | P1,P2,P3 | P2 | 1/5,1,1/3 | 0.1062 | 3.039 | 0.019 | 0.037 | Pass |
| C1 sub-criteria layer | C1-P | P1,P2,P3 | P3 | 1/3,3,1 | 0.2605 | 3.039 | 0.019 | 0.037 | Pass |
| C2 sub-criteria layer | C2-P | P4,P5,P6 | P4 | 1,1/3,1/5 | 0.1062 | 3.039 | 0.019 | 0.037 | Pass |
| C2 sub-criteria layer | C2-P | P4,P5,P6 | P5 | 3,1,1/3 | 0.2605 | 3.039 | 0.019 | 0.037 | Pass |
| C2 sub-criteria layer | C2-P | P4,P5,P6 | P6 | 5,3,1 | 0.6333 | 3.039 | 0.019 | 0.037 | Pass |
| C3 sub-criteria layer | C3-P | P7,P8,P9 | P7 | 1,3,2 | 0.5390 | 3.009 | 0.005 | 0.009 | Pass |
| C3 sub-criteria layer | C3-P | P7,P8,P9 | P8 | 1/3,1,1/2 | 0.1638 | 3.082 | 0.041 | 0.079 | Pass |
| C3 sub-criteria layer | C3-P | P7,P8,P9 | P9 | 1/2,2,1 | 0.2973 | 3.082 | 0.041 | 0.079 | Pass |
| C4 sub-criteria layer | C4-P | P10,P11,P12 | P10 | 1,3,1/3 | 0.2605 | 3.039 | 0.019 | 0.037 | Pass |
| C4 sub-criteria layer | C4-P | P10,P11,P12 | P11 | 1/3,1,1/5 | 0.1062 | 3.039 | 0.019 | 0.037 | Pass |
| C4 sub-criteria layer | C4-P | P10,P11,P12 | P12 | 3,5,1 | 0.6333 | 3.039 | 0.019 | 0.037 | Pass |

### Calculation of Comprehensive Evaluation Value and Grade Classification

After obtaining the criterion-level weights and indicator-level local weights, the global weight of each indicator relative to the objective level was calculated as follows:

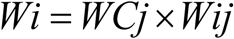

In the formula, *Wi* denotes the global weight of the *i*-th indicator relative to the objective layer; *WCj* denotes the weight of the *j*-th criterion relative to the objective layer; and *Wij* denotes the local weight of the *i*-th indicator relative to the corresponding criterion layer. The global weights of the *H. mutabilis* hierarchy are presented in Table 5.

**Table 5.** The overall priority of the criterion layer P with respect to the goal layer A.

| **Criterion C** | **C weight to A** | **Indicator P** | **P weight to C** | **P weight to A** |
|---|---|---|---|---|
| C1 | 0.4168 | P1 | 0.6333 | 0.2640 |
|  |  | P2 | 0.1062 | 0.0442 |
|  |  | P3 | 0.2605 | 0.1086 |
| C2 | 0.2695 | P4 | 0.1062 | 0.0286 |
|  |  | P5 | 0.2605 | 0.0702 |
|  |  | P6 | 0.6333 | 0.1707 |
| C3 | 0.1209 | P7 | 0.5390 | 0.0652 |
|  |  | P8 | 0.1638 | 0.0198 |
|  |  | P9 | 0.2973 | 0.0360 |
| C4 | 0.1928 | P10 | 0.2605 | 0.0502 |
|  |  | P11 | 0.1062 | 0.0205 |
|  |  | P12 | 0.6333 | 0.1221 |

Based on the measured values and scoring criteria, each indicator was classified into three levels—5, 3, and 1 points—with 2 and 4 points as intermediate scores between adjacent levels (Table 6). The score for each cultivar on each indicator is denoted as Pi; the composite score is calculated using the total weighting Wi for the indicators. The formula for the composite score is

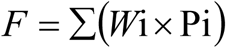

**Table 6.** Grading Standards for Evaluation Indicators of *Hibiscus mutabilis* Food.

| Indicator | Score 5 | Score 3 | Score 1 |
|---|---|---|---|
| P1 Heavy metal contamination risk | ≤ 64.15 µg/kg | 96.88–129.61 µg/kg | > 162.34 µg/kg |
| P2 Microbiological contamination risk | Total count < 1×10<sup>6</sup> CFU/g; E. coli < 10 | Total count 5×10<sup>6</sup>–1×10<sup>7</sup> CFU/g; E. coli 100–1×10<sup>3</sup> | Total count > 5×10<sup>7</sup> CFU/g; E. coli > 1×10<sup>6</sup> |
| P3 Endogenous toxin risk (free gossypol) | ≤ 20 mg/kg | 22–27 mg/kg | > 30 mg/kg |
| P4 Total flavonoid content | > 0.040 g/100 g | 0.025–0.030 g/100 g | < 0.020 g/100 g |
| P5 Total amino acid content | ≥ 12.00 mg/g | 8.50 ≤ T < 10.00 mg/g | < 7.50 mg/g |
| P6 Total mineral element content | ≥ 34 000 mg/kg | 30 000–32 000 mg/kg | < 28 000 mg/kg |
| P7 Fiber coarseness | Not rough | Clearly rough | Strongly rough |
| P8 Lingering astringency (tannin, g/100 g) | < 3.50 | 4.00–4.79 | ≥ 5.60 |
| P9 Mouthfeel smoothness | Honey-smooth, no discomfort | Slight roughness | Obvious throat irritation |
| P10 Yield per unit area | ≥ 975 kg/ha | 525–749.85 kg/ha | < 375 kg/ha |
| P11 Drying yield (dry/fresh ratio) | ≥ 0.20 | 0.13–0.149 | < 0.115 |
| P12 Storability (browning degree at the same time) | < 30% browning | 30%–70% browning | ≥ 90% browning |

where F is the comprehensive evaluation value of a given cultivar; Wi is the total weight of indicator i relative to the target layer; Pi is the score of the cultivar on indicator i; and n is the number of evaluation indicators (n = 12 in this study). Cultivars with F ≥ 3.5 were classified as Grade I (high edible quality); 3.0 ≤ F < 3.5 as Grade II (moderate); and F < 3.0 as Grade III (limited).

#### Data Processing and Statistical Analysis

All measurements were performed in triplicate, and the results are expressed as Mean ± SD. Raw data were processed in Excel 2021 to compute the mean, SD, and composite score for each parameter; each parameter was then converted to a 1–5 score according to the predefined scoring criteria. AHP weights were computed and judgement matrix consistency was verified using SPSSAU. Differences among cultivars in measured parameters were assessed by one-way analysis of variance (ANOVA), with significance defined as *P* < 0.05. Bar charts and heatmaps were generated using Origin 2022 software to visualize the key evaluation indicators across cultivars and the corresponding indicator-level scores.

## Results

### Measured Values of Comprehensive Evaluation Indicators

Notable differences among *H. mutabilis* cultivars were observed in safety, nutritional value, and bioactive compound profiles (Fig 2).After screening against safety standards, heavy metal analysis revealed distinct accumulation patterns among the 11 cultivars. Specifically, ‘Jinqiuhong’, ‘Danbanhong’, ‘Jinbiyu’, and ‘Jinxiuzi’ accumulated relatively higher levels of certain heavy metals, whereas ‘Jinqiusong’ and ‘Bairi Huacai’ showed lower levels, indicating inter-cultivar variation in measured heavy metal concentrations. *Staphylococcus aureus* was not detected in any of the 11 cultivars. However, other microbial indicators varied among cultivars. Variations in total flavonoid count, mould count, and coliform count were observed among cultivars; specifically, ‘Bairi Huacai’, ‘Chongbanbai’, and ‘Mudanfen’ showed relatively high levels of certain microbial indicators, whereas ‘Jinxiuzi’ presented a comparatively low overall contamination risk. Free gossypol was detected in only four cultivars, with the highest concentration observed in ‘Mudanfen’, followed by ‘Jinbiyu’, ‘Caixia’, and ‘Danbanhong’. Concentrations in the remaining cultivars were below the detection limit, indicating variation in free gossypol concentrations among cultivars.

**Fig 2.**
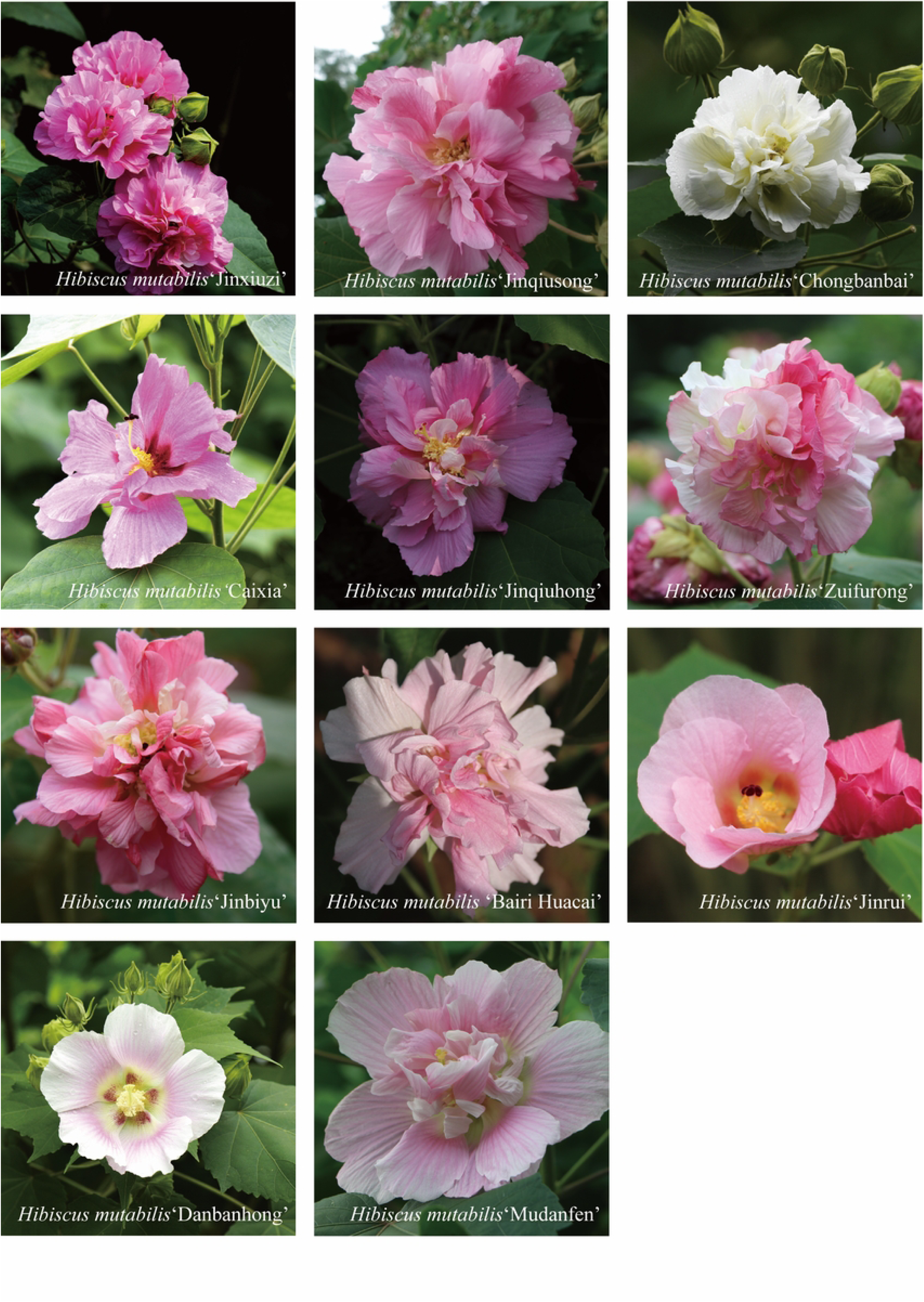
Comparison of Safety, Nutritional, and Active Substance Indicators among Different *Hibiscus mutabilis* Cultivars. Note:(A) Heatmap of heavy metal elements. (B) Heatmap of microbial contamination indicators. (C) Gossypol-related toxicity. (D) Total flavonoid content. Values are Means ± SD (n = 3). Different lowercase letters denote significant differences among cultivars at *P* < 0.05 (one-way ANOVA). “ND” indicates not detected (below the detection limit).

Among nutritional and active substance, notable variations were observed in total flavonoid, amino acid, and mineral element concentrations across cultivars. ‘Jinxiuzi’ exhibited the highest total flavonoid content, followed by ‘Zuifurong’. For amino acids, ‘Jinxiuzi’, ‘Jinqiusong’, and ‘Chongbanbai’ ranked highest. Regarding mineral elements, ‘Jinqiuhong’, ‘Danbanhong’, ‘Caixia’, and ‘Mudanfen’ showed comparatively higher total mineral levels, whereas ‘Bairi Huacai’ and ‘Jinqiusong’ were relatively lower.Among sensory parameters, total tannin content varied considerably across cultivars. ‘Bairi Huacai’ exhibited the highest total tannin content, followed by ‘Caixia’ and ‘Chongbanbai’, indicating a relatively high risk of lingering astringency. Conversely, ‘Danbanhong’, ‘Jinqiusong’, and ‘Jinrui’ showed lower total tannin content, suggesting relatively better sensory palatability. Yield per unit area results showed that ‘Bairi Huacai’ achieved the highest fresh and dried flower yields (kg/ha). ‘Chongbanbai’, ‘Jinxiuzi’, ‘Zuifurong’, and ‘Jinqiuhong’ ranked second, whereas ‘Danbanhong’, ‘Mudanfen’, and ‘Jinrui’ showed relatively lower yields (Fig 3).

**Fig 3.**
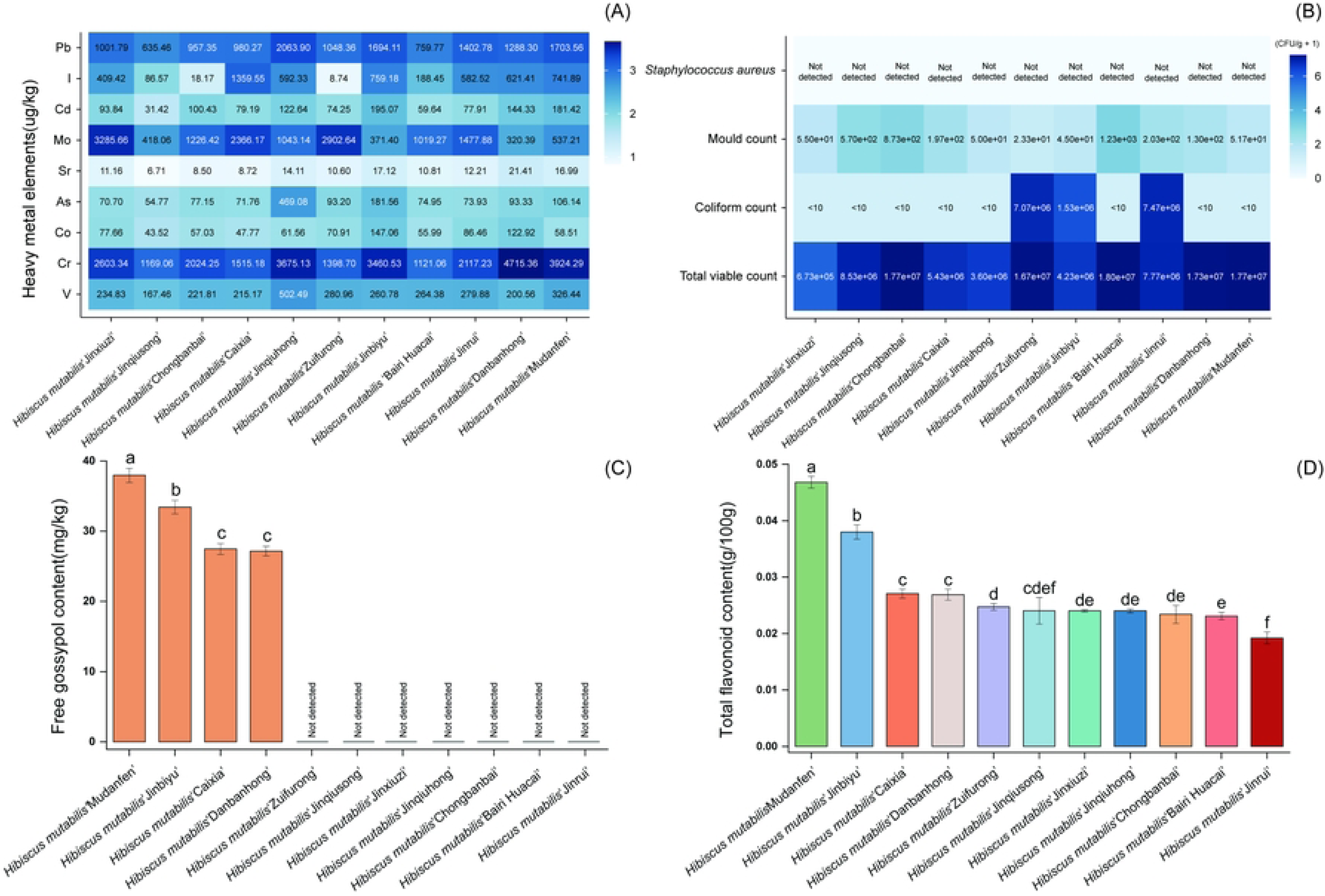
Comparison of Nutritional, Active Substances, and Yields among Different *Hibiscus mutabilis* Cultivars. Note: (A) Heatmap of amino acid profiles. (B) Heatmap of mineral element concentrations. (C) Total tannin content. (D) Yield per unit area of dried flowers and fresh flowers. Values are means ± SD (n = 3). Different lowercase letters denote significant differences among cultivars at *P* < 0.05.

### Indicator-Layer Scores of Different Cultivars

Based on the measured parameters and sensory evaluation results, the performance of the 11 cultivars across all 12 indicators was converted to 1–5 scores according to the predefined scoring criteria; the results are presented in Fig 4.Based on the scoring results, ‘Jinqiusong’, ‘Bairi Huacai’, and ‘Jinbiyu’ performed best on the heavy metal contamination risk, whereas ‘Jinxiuzi’ and ‘Jinqiuhong’ scored highest on the microbial contamination risk index. For gossypol-related toxicity risk, ‘Zuifurong’, ‘Jinqiusong’, ‘Jinqiuhong’, ‘Chongbanbai’, ‘Bairi Huacai’, and ‘Jinrui’ ranked highest. Among nutritional and active substance, ‘Jinxiuzi’ led in total flavonoid and total amino acid content, while ‘Danbanhong’, ‘Jinqiuhong’, and ‘Mudanfen’ ranked highest in mineral element concentrations. For sensory quality, most cultivars scored well on fibre coarseness, whereas marked differences were observed in lingering astringency and mouthfeel smoothness. Regarding raw material supply characteristics, ‘Bairi Huacai’ ranked highest in yield per unit area, ‘Chongbanbai’ in storability, and ‘Danbanhong’ in drying yield.

**Fig 4.**
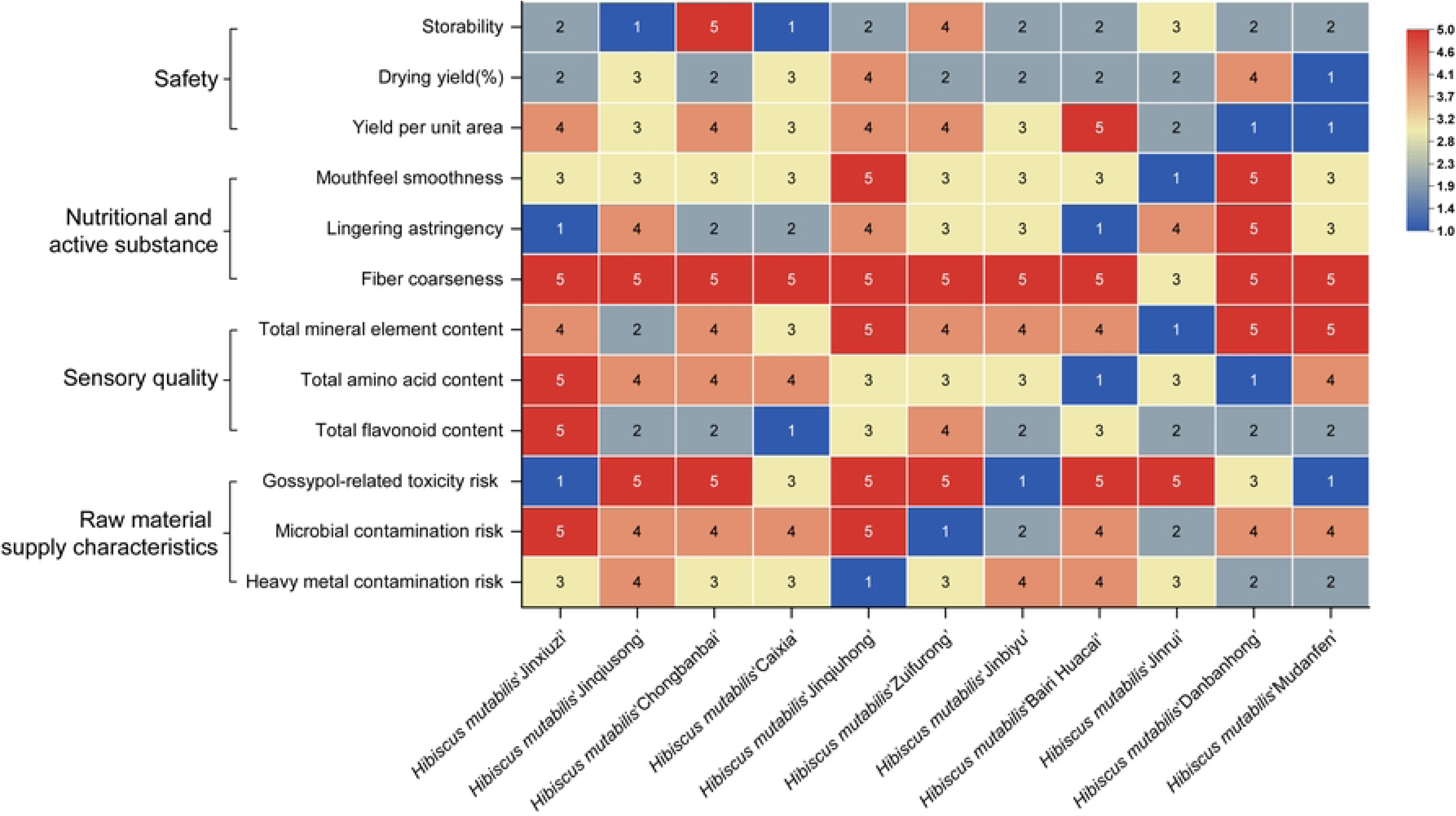
Scores of Criterion Layer Indicators among Different *Hibiscus mutabilis* Cultivars. Note: Comprehensive quality evaluation of 11 *Hibiscus mutabilis* cultivars based on raw material supply characteristics, sensory quality, nutritional and active substance, and safety-related indicators. Each indicator was evaluated using a five-point scoring system, with higher scores indicating better quality performance. Undesirable attributes, including lingering astringency, fiber coarseness, endogenous toxin risk, microbial contamination risk, and heavy metal contamination risk, were scored so that higher scores indicate less pronounced undesirable characteristics or lower risks. Colors range from blue (1, poorest performance) to red (5, best performance)

### Comprehensive Evaluation Values and Grade Classification

Based on the indicator scores and their global weights relative to the objective layer, comprehensive evaluation values were calculated for the 11 cultivars, ranked, and assigned to grade levels, as presented in Table 7. For example, ‘Bairi Huacai’ achieved a weighted comprehensive score of 3.6043 across the 12 indicators, corresponding to Grade I.

**Table 7.** Comprehensive evaluation scores and grading of 11 *Hibiscus mutabilis* Cultivars.

| **Rank** | **Cultivar** | **Final score** | **Grade** |
|---|---|---|---|
| 1 | 'Bairi Huacai' | 3.6043 | I |
| 2 | 'Chongbanbai' | 3.5533 | I |
| 3 | 'Jinqiusong' | 3.5055 | I |
| 4 | 'Jinqiuhong' | 3.4930 | II |
| 5 | 'Zuifurong' | 3.4069 | II |
| 6 | 'Danbanhong' | 3.3178 | II |
| 7 | 'Jinxiuzi' | 3.2380 | II |
| 8 | 'Jinbiyu' | 3.1325 | II |
| 9 | 'Caixia' | 3.1269 | II |
| 10 | 'Mudanfen' | 2.7113 | III |
| 11 | 'Jinrui' | 2.5785 | III |

As presented in Table 7, the comprehensive evaluation scores of the 11 cultivars ranged from 2.5785 to 3.6043. ‘Bairi Huacai’ achieved the highest score (3.6043), followed by ‘Chongbanbai’ (3.5533) and ‘Jinqiusong’ (3.5055). Based on the grading criteria, three cultivars were classified as Grade I, representing 27.27% of the samples: ‘Bairi Huacai’, ‘Chongbanbai’, and ‘Jinqiusong’, indicating superior edible quality. Six cultivars were classified as Grade II (54.55%): ‘Jinqiuhong’, ‘Zuifurong’, ‘Danbanhong’, ‘Jinxiuzi’, ‘Jinbiyu’, and ‘Caixia’, representing an intermediate overall performance. The remaining two cultivars, ‘Mudanfen’ and ‘Jinrui’, were classified as Grade III (18.18%), with relatively lower overall scores.

## Discussion

### Analysis of Evaluation Indicator Weights

As shown by the criterion layer weights relative to the goal layer, safety received the highest weight, followed by nutritional and active substance, raw material supply characteristics, and sensory quality. This confirms that safety is the primary consideration when transitioning *Hibiscus mutabilis* L. from ornamental plants to food resources[24–25].

At the indicator layer, the global weights of measured heavy metal concentrations, mineral element concentrations, and gossypol-related toxicity risk have the greatest influence on the comprehensive evaluation results, indicating that safety, together with nutritional and active substance, jointly determines the priority of *H. mutabilis* flowers for food development[26–27]. By comparison, sensory indicators such as fibre coarseness and mouthfeel smoothness received relatively lower weights; nevertheless, given their influence on consumer acceptance and product palatability, they should not be neglected in future product development[28–29].

The AHP-derived weights primarily reflect the relative importance assigned by experts and are inherently influenced by the composition of the expert panel, the construction of the judgement matrix, and the definition of evaluation objectives[30–31]. Accordingly, the evaluation model established in this study is best suited for preliminary screening of *H. mutabilis* cultivars; further validation should incorporate empirical measurement data, food safety standards, and product processing trials.

### Analysis of Comprehensive Evaluation Results

Based on the comprehensive evaluation scores, the 11 cultivars were classified into three grades. Three cultivars (27.27%) were assigned to Grade I: ‘Bairi Huacai’, ‘Chongbanbai’, and ‘Jinqiusong’. Six cultivars (54.55%) were assigned to Grade II: ‘Jinqiuhong’, ‘Zuifurong’, ‘Danbanhong’, ‘Jinxiuzi’, ‘Jinbiyu’, and ‘Caixia’. The remaining two cultivars (18.18%) were assigned to Grade III: ‘Mudanfen’ and ‘Jinrui’.

Of the 11 cultivars, ‘Bairi Huacai’ (3.6043)achieved the highest overall score under the present weighting scheme. However, its high total tannin content suggests a potential limitation related to astringency that warrants further sensory evaluation. ‘Chongbanbai’ ranked second (3.5533), with excellent safety and several nutritional and active substance indicators; however, it was relatively weak in storability. If used for food processing or fresh produce development, complementary post-harvest preservation and short-term storage technologies should be prioritized. ‘Jinqiusong’ scored 3.5055, meeting Grade I criteria but with a comparatively lower overall score.

Grade II cultivars achieved composite scores between 3.0 and 3.5, indicating intermediate performance under the present evaluation framework. These cultivars exhibited strengths in certain indicators but limitations in others, suggesting that their suitability for food development may depend on specific processing requirements and intended applications.

